# Horizontal gene cluster transfer increased hallucinogenic mushroom diversity

**DOI:** 10.1101/176347

**Authors:** Hannah T. Reynolds, Vinod Vijayakumar, Emile Gluck-Thaler, Hailee Brynn Korotkin, Patrick Brandon Matheny, Jason Christopher Slot

## Abstract

Secondary metabolites are heterogeneous natural products that often mediate interactions between species. The tryptophan-derived secondary metabolite, psilocin, is a serotonin receptor agonist that induces altered states of consciousness. A phylogenetically disjunct group of mushroom-forming fungi in the Agaricales produce the psilocin prodrug, psilocybin. Spotty phylogenetic distributions of fungal compounds are sometimes explained by horizontal transfer of metabolic gene clusters among unrelated fungi with overlapping niches. We report the discovery of a psilocybin gene cluster in three hallucinogenic mushroom genomes, and evidence for its horizontal transfer between fungal lineages. Patterns of gene distribution and transmission suggest that psilocybin provides a fitness advantage in the dung and late wood-decay niches, which may be reservoirs of fungal indole-based metabolites that alter behavior of mycophagous and wood-eating invertebrates. These hallucinogenic mushroom genomes will serve as models in neurochemical ecology, advancing the prospecting and synthetic biology of novel neuropharmaceuticals.

---

Secondary metabolites (SMs) are small molecules that are widely employed in defense, competition, and signaling among organisms (Raguso et al. 2015). Due to their physiological activities, SMs have been adopted by both ancient and modern human societies as medical, spiritual, or recreational drugs. Psilocin is a psychoactive agonist of the serotonin (5-hydroxytryptamine, **5-HT**) -2a receptor (Halberstadt and Geyer 2011) and is produced as the phosphorylated prodrug psilocybin by a restricted number of distantly related mushroom forming families of the Agaricales (Bolbitiaceae, Inocybaceae, Hymenogastraceae, Pluteaceae) (Allen 2010 May 19; Dinis-Oliveira 2017). Hallucinogenic mushrooms have a long history of religious use, particularly in Mesoamerica, and were a catalyst of cultural revolution in the West in the mid 20th century (Nyberg 1992; Letcher 2006). Psilocybin was structurally described and synthesized in 1958 by Albert Hoffman (Hofmann et al. 1958), and a biosynthetic pathway (Fig. 1B) was later proposed based on the transformation of precursor molecules by *Psilocybe cubensis* (Agurell and Nilsson 1968). However, prohibition since the 1970s (21 U.S. Code § 812 -Schedules of controlled substances) has limited advances in psilocybin genetics, ecology, and evolution. There has been a recent resurgence of the hallucinogen research in the clinical setting. Brain state imaging studies of psilocin exposure have identified changes in neural activity and interconnectivity that underlie subjective experiences, and therapeutic trials have investigated psilocybin’s potential for treating major depression and addictive disorders (Griffiths et al. 2011; Carhart-Harris et al. 2012; Petri et al. 2014; Carhart-Harris et al. 2016; Johnson et al. 2017). While the ecological roles of psilocybin, like most SMs, remain unknown, psilocin’s mechanism of action suggests metazoans may be its principal targets.

**Figure 1.**
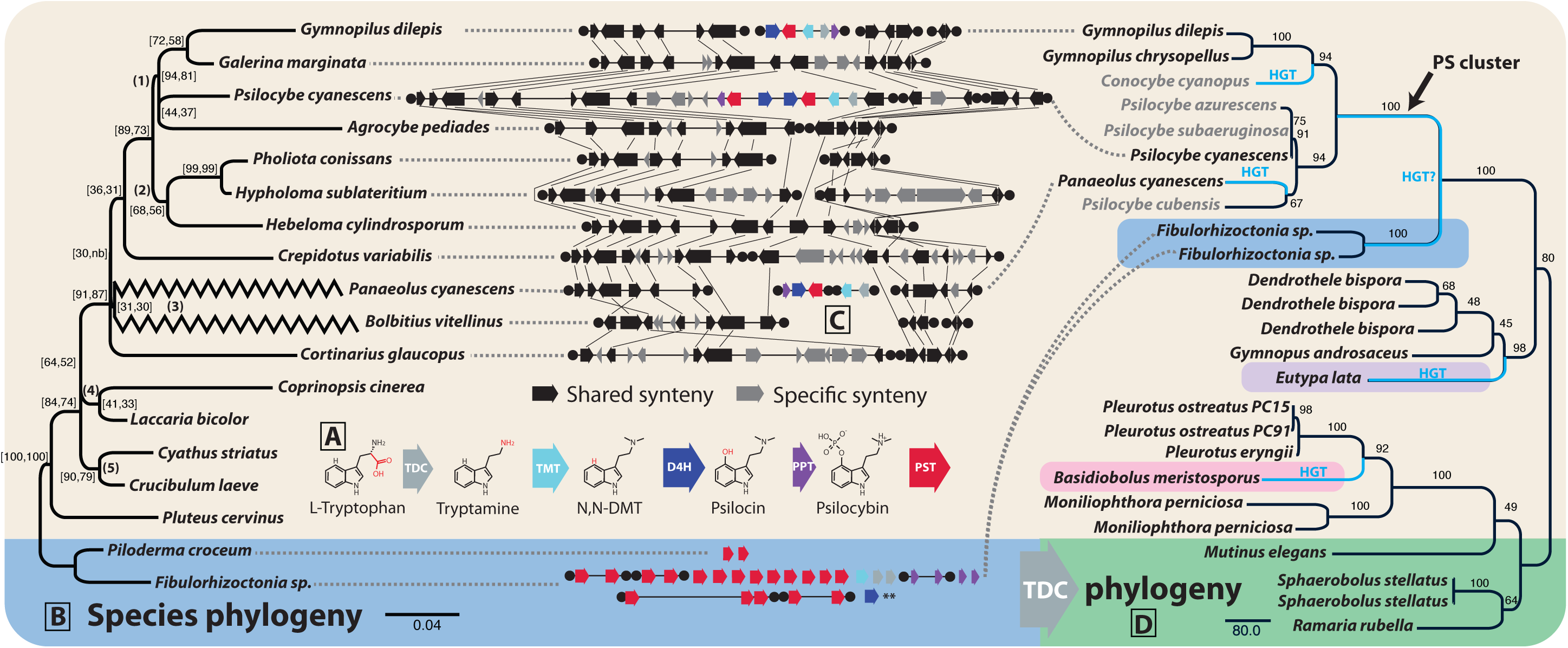
Psilocybin evolution. A. The PS cluster consists of tryptophan decarboxylase (TDC), 1-2 P450 monooxygenases (D4H), methyltransferase (TMT), phosphotransferase (PPT), and 1-2 MFS transporters (PST). B. Phylogenomic tree of Agaricales (tan) with Atheliales (blue) outgroup. Support values = (internode certainty, tree certainty). Clades 1-5 are as in Fig. 2B. C. PS locus synteny relative to *Ps. cyanescens* scaffold 5617 and *G. marginata* scaffold 9. D. RAxML phylogeny of TDC indicating putative HT branches; *Eutypa lata* is in Xylariales (Ascomycota, lavendar), an order correlated with absence of termites in coarse woody debris (Kirker et al. 2012), with members that produce a white rot of wood. Entomophthoromycotina sp. 1, rose) is commonly associated with amphibian dung and arthropods. Grey taxon names = PCR sequences, black = whole genome. Support is percent of 100 ML bootstraps. **54 similar D4H homologs not shown.

A common feature of fungal SM biosynthesis is the organization of most or all required anabolic, transport, and regulatory elements in gene clusters (GCs). GCs are often discontinuously distributed among fungal taxa, partly due to horizontal transfer (HT) among species with overlapping ecological niches (Gluck-Thaler and Slot 2015). The sparse phylogenetic distribution of psilocybin, coupled with the requirement for multiple enzymatic steps for its biosynthesis (tryptophan-decarboxylation, N-methylation, indole-4-hydroxylation, and O-phosphorylation), suggest the psilocybin pathway might have dispersed via horizontal GC transfer, and therefore the genetic mechanism for psilocybin biosynthesis might be identified in searches for GCs with a common phylogenetic history and distribution among psilocybin producing (PS+) mushrooms. The pattern of HT events may further suggest ecological pressures that have driven the pathway’s persistence (Baquero 2004).

We identified candidate psilocybin genes by sequencing three diverse PS+ mushroom monokaryon genomes --*Psilocybe cyanescens*, *Panaeolus* (=*Copelandia*) *cyanescens*, and *Gymnopilus dilepis* (Table 1), and comparing them to three related mushrooms not known to produce psilocybin (PS-): *Galerina marginata*, Agaricales sp.9, and *Hypholoma sublateritium*. Of 37 gene homolog groups (HGs) consistent with a PS+ distribution among these taxa, only five were clustered, all in PS+ genomes (Fig. S1-pipeline and HGs). We retroactively designated *Gy. chrysopellus*, potentially PS+ because it possesses a cluster identical to *Gy. dilepis*, which is not a surprising oversight given inconsistent identifications, and geographical variation among *Gymnopilus* spp. phenotypes. Predicted functions of these five genes were also consistent with psilocybin biosynthesis and metabolite transport, and were putatively designated tryptophan decarboxylase (TDC), tryptamine N-methyltransferase (TMT), dimethyltryptamine-4-hydroxylase (D4H), psilocin phosphotransferase (PPT), and psilocybin transporter (PST). As SM GCs are infrequently identified in Basidiomycota compared with Ascomycota, this is a notable discovery (Quin et al. 2014).

**Table 1.** Genome assembly and annotation of psilocybin-producing mushrooms.

|  | length (nt) | scaffolds | contigs | av. depth of coverage (x) | N50 (nt) | Complete BUSCOs (%) | total proteins | decarboxylases / PSD <sup>a</sup> | P450's<sup>a</sup> | methyltransferases / DUF890 domain-proteins<sup>a</sup> | kinases / phosphotransferases / OG term 0PNAW<sup>a</sup> | MFS<sup>a</sup> |
| --- | --- | --- | --- | --- | --- | --- | --- | --- | --- | --- | --- | --- |
| G. dilepis<sup>b</sup><br>TENN071165 | 47,177,497 | 8,423 | 10,681 | 16.5 | 33,540 | 73.43 | 16,257 | 28 / 9 | 151 | 89 / 4 | 275 / 29 / 1 | 37 |
| Pa. cyanescens | 44,965,162 | 9,521 | 11,850 | 25.7 | 32,751 | 75.66 | 13,420 | 28 / 8 | 148 | 91 / 2 | 267 / 16 / 2 | 34 |
| Ps. cyanescens | 53,483,841 | 18,721 | 38,006 | 44.7 | 46,250 | 72.18 | 15,973 | 38 / 17 | 178 | 102 / 2 | 298 / 23 / 1 | 44 |

<sup>a</sup>Functional category of PS genes as annotated in Eggnog. <sup>b</sup>*G. dilepis* (= *G. aeruginosus* sensu L.R. Hesler) was isolated from oak sawdust in Knoxville, Tennessee 4-Oct-2013. *Pa. cyanescens* and *Ps. cyanescens* basidiospores were supplied by The Spore Works, Knoxville.

To confirm GC function, we profiled the enzymology of heterologously expressed TDC and PPT, and assayed by LC-MS/MS analyses. We determined that TDC, the first committed step in the reaction and the only one not producing a drug-scheduled compound, has specific decarboxylase activity on tryptophan. TDC reactions produced tryptamine, identified at the characteristic m/z 144.1 [M+H]+, (Fig. S2, Supplemental data1). TDC did not decarboxylate phenylalanine, tyrosine or 5-hydroxy-L-tryptophan (5-HTP) under the same conditions. We note that TDC is similar to type II phosphatidylserine decarboxylases (PSDs), but has no significant sequence similarity with a pyridoxal-5’-phosphate-dependent decarboxylase recently characterized in *Ceriporiopsis subvermispora* as specific for L-tryptophan and 5-HTP (Kalb et al. 2016). A unique GGSS sequence in a conserved C-terminal motif (Fig. S3), suggests tryptophan decarboxylation is a previously unknown derived function among PSDs (Wriessnegger et al. 2009; Choi et al. 2015). We detected no activity of PPT on 5-HT or 4-Hydroxyindole (4-HI) as alternatives to the psilocin substrate, possibly due to requirements for the 4-hydroxyl and the methylated amine groups of psilocin, but further characterization of PPT and other enzymes was prevented by the regulatory status of substrates and products.

Phylogenetic analyses of PS homologs from a local database of 618 fungal proteomes yielded congruent gene tree topologies with respect to PS+ taxa, and clades of clustered PS genes from all gene trees excluded the PS-taxa in the database, suggesting the clustered genes are coordinately-inherited (Fig. 1, Fig. S4 A-F).The gene trees also suggest HT of the cluster from *Psilocybe* to *Panaeolus* and HT of most PS genes between Atheliaceae and Agaricaceae when compared to a phylogenomic tree of related Agaricales (Fig. 1). The direction of the latter HT is ambiguous, and not strongly supported by all five genes. Analyses with TDC and PPT amplicon sequences retrieved by degenerate PCR of unsequenced *Psilocybe* and *Conocybe* genomes (Supplemental data1) suggest the dung fungus *Ps. cubensis* vertically inherited the cluster, and *Pa. cyanescens* acquired the cluster from *Psilocybe* sp., and possibly from a dung-associated lineage. Alternative hypotheses of vertical inheritance in these lineages were rejected; exclusion of *Pa. cyanescens* and *C. cyanopus* (AU test, p = 0.004) or *Pa. cyanescens* alone (p = 0.036) were significantly worse constrained topologies (Supplemental data1). Furthermore, a TDC gene tree-species tree reconciliation model allowing duplication, HT, and loss (6 events: D=1, HT=3, L=2) is more parsimonious than a model that only allows duplication and loss (28 events: D=3, L=25)(Fig. S5). PS gene orthologs were not detected in *Ps. fuscofulva*, a PS-species representing an early branch in *Psilocybe* diversification (Borovička et al. 2015). Conservation of synteny flanking the *Ps. cyanescens* PS cluster (Fig. 1) suggests it may have been recently acquired in *Psilocybe* as well, or lost as a unit in close relatives. A genome wide scan did not identify additional HT genes or clusters between *Psilocybe* and *Panaeolus* (Fig. S6). HT is comparatively rare in the Basidiomycetes, suggesting the transfer of the PS cluster may have special significance (Wisecaver et al. 2014), and is to our knowledge, the first report of HT of a SM GC between lineages of mushroom-forming fungi (Agaricomycotina).

Recent studies suggest that ecology can select for both genome content (Ma et al. 2010; de Jonge et al. 2013) and organization in eukaryotes through both vertical and horizontal patterns of inheritance (Holliday et al. 2015; Kakioka et al. 2015). Ordination of 10,998 HGs identified two principal components (PCs) that describe 22% of the variation in gene content among 16 Agaricales genomes (Fig. 2B). Discrimination of genome composition along PC1 appears to reflect phylogenetic differences, while discrimination along PC2 parallels ecological differences between plant mutualists and other fungi. However, PC2 does not discriminate between dung and wood-decay fungi. The functions of HGs most associated with each PC are consistent with this interpretation. All eight metabolism-related processes in the COG classification system are overrepresented in PC2, but only one is overrepresented in PC1 HGs (Supplemental data1). The grouping of several divergent lineages of wood and dung decay fungi to the exclusion of close ectomycorrhizal relatives along PC2 may reflect similar selective pressures in the decayed wood and dung environments, from recalcitrant plant polymers like lignin, and invertebrate predation (Rouland-Lefèvre 2000). However, a small number of HGs exclusive to either wood or dung associated fungi (Fig. S7, Supplemental data1) are consistent with ecological specialization within each guild. Wood-specific genes include functions in lignin degradation (e.g., peroxidase, isoamyl alcohol oxidase) and carbohydrate transport, while dung-specific genes have functions in bacterial cell wall degradation (e.g., lysozyme), hemicellulose degradation (e.g., Endo-1,4-beta-xylanase, Alpha-L-arabinofuranosidase), and inorganic phosphate transport. Niche-specific genes are largely consistent with vertical inheritance; however, analyses support HT of a single ferric-reductase-like gene (pfam01794, pfam00175) likely involved in iron uptake, to Coprinopsis and Panaeolus from dung-associated Ascomycota (Supplemental data).

**Figure 2.**
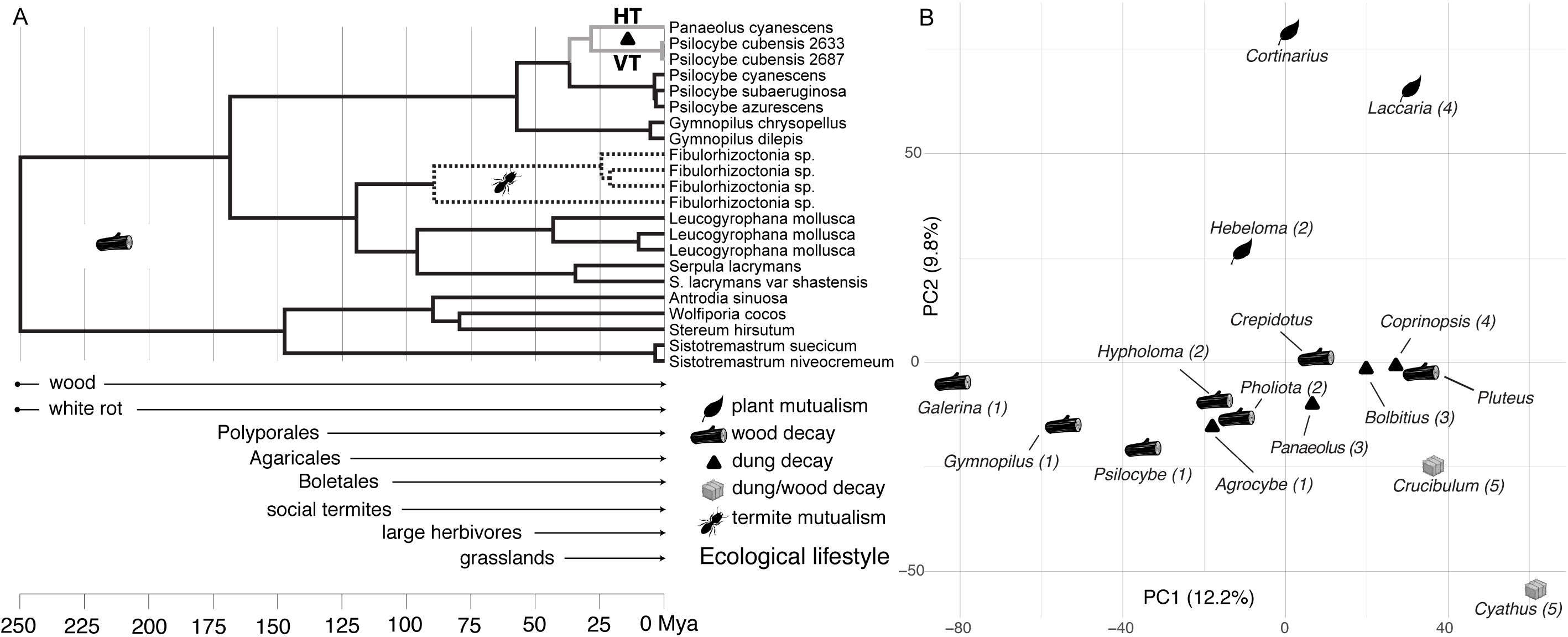
Patterns of ecological diversification of PS genes and Agaricales genomes. A. Ultrametric representation of PPT phylogeny, with root age hypothetically set to align ecological transitions with Earth history events. HT = horizontal transfer, VT = vertical transmission. B. Ordination of Agaricales genome content. Numerals correspond to clades in Fig. 1A.

In addition to similar ecological pressures, similar genome content among wood and dung-decaying fungi may also reflect the ecological diversification of Agaricomycetes that accompanied major geological transformations (Fig. 2B). For example, the emergence of true wood opened a massive saprotrophy niche space in the upper Devonian (380 Mya), in which the Agaricomycetes diversified with the aid of key enzymatic innovations (Floudas et al. 2012). The subsequent radiation of herbivorous megafauna during the Eocene approximately 50 MYA (MacFadden 2000) and the spread of grasslands 40 MYA (Retallack 2001) expanded the mammalian dung niche space in which invertebrates and fungi competed. These changes parallel the repeated emergence of dung-specialization from plant-decay ancestors in the radiation of *Psilocybe* and other Agaricales lineages (Ramirez-Cruz et al. 2013; Tóth et al. 2013). Late stage wood decay fungi like *Psilocybe* spp. likely harbor genetic exaptations for lignin tolerance/degradation, and competition with invertebrates and prokaryotes, and acquisition of particularly adaptive functions by other fungi (e.g. *Panaeolus*) through HT may have facilitated additional transitions to dung saprotrophy.

The evolution of PS genes suggest they originally served roles in the wood-decay niche, and more recently emerged through both vertical and horizontal transfer in dung-decay fungi (Fig. 2). HT and retention of PS clusters are evidence of selection on the PS pathway in the recipient lineage, as SM clusters are inherently unstable in fungal genomes (Reynolds et al. 2017 Apr 28). Psilocybin neurological activity, coupled with HT and retention in lineages that colonize dung and/or decayed wood, which are rich in both mycophagous and competitor invertebrates (Rouland-Lefevre 2000), suggest that psilocybin may be a modulator of insect behavior. Psilocybin and/or the related aeruginascin have also been identified in the lichenized agaric, *Dictyonema huaorani*, and in the ectomycorrhizal genus *Inocybe* (Kosentka et al. 2013; Schmull et al. 2014). PS distribution in *Inocybe* is complementary to that of the acetylcholine mimic, muscarine, which could suggest alternative strategies and pressures to manipulate animal behavior beyond the dung and wood decay niches. Neurotransmitter mimics may provide advantages to fungi by interfering with the behavior of invertebrate competitors for woody resources (Hunt et al. 2007), especially social insects, like termites, which emerged ∼ 137 Mya, because they rely on the coordinated activities of multiple castes (Genise 2017). It is thus intriguing that D4H and PST have experienced massive gene family expansion through duplication in *Fibulorhizoctonia* sp., which produce termite egg-mimicking sclerotia in an ancient mutualistic relationship with *Reticulitermes* termites (Matsuura 2005). While neurotransmitter agonists are not known to mediate this symbiosis, insect predatory fungi (i.e. *Cordyceps* spp.) use neurotransmitter analogs to influence the behavior of infected insects (de Bekker et al. 2014), and a number of repellents and toxins in wood-decay fungi inhibit xylophagy and mycophagy by termites (Rouland-Lefèvre 2000).

The identification of genes underlying PS biosynthesis is an important advance in the field of neurochemical ecology, with both social and medical applications. The sequences of the first *Psilocybe* and *Panaeolus* genomes presented here will be important resources for the prospecting of novel neurotropic natural products (Rutledge and Challis 2015). The discovery that a psilocybin cluster has been horizontally transferred and subsequently maintained among the invertebrate-challenged environments of dung and late wood-decay suggests these niches may be reservoirs not only of new antibiotics (Bills et al. 2013 Aug 23), but also novel neuroactive pharmaceuticals.

## Acknowledgements

The authors thank Jan Borovička and Paul Stamets for providing additional cultures used in this study. We also thank Michael Kelly, Ma Elena Hernandez, Stephen Opiyo, and Tea Meulia, OARDC Molecular and Cellular Imaging Center for technical support. Computation was performed using The Ohio Supercomputer Center resources. Genome assemblies are deposited in GenBank under accessions SAMN07166449, SAMN07169033, and SAMN07169108.

